# Comparing three types of dietary samples for prey DNA decay in an insect generalist predator

**DOI:** 10.1101/098806

**Authors:** Stefaniya Kamenova, Rebecca Mayer, Oskar R. Rubbmark, Eric Coissac, Manuel Plantegenest, Michael Traugott

## Abstract

The rapidly growing field of molecular diet analysis is becoming increasingly popular among ecologists, especially when investigating methodologically challenging groups such as invertebrate generalist predators. Prey DNA detection success is known to be affected by multiple factors, however the type of dietary sample has rarely been considered. Here, we address this knowledge gap by comparing prey DNA detection success from three types of dietary samples. In a controlled feeding experiment, using the carabid beetle *Pterostichus melanarius* as a model predator, we collected regurgitates, feces and whole consumers (including their gut contents) at different time points post-feeding. All dietary samples were analyzed using multiplex PCR targeting three different length DNA fragments (128 bp, 332 bp and 612 bp). Our results show that both the type of dietary sample and the size of the DNA fragment contribute to a significant part of the variation found in the detectability of prey DNA. Specifically, we observed that in both regurgitates and whole consumers prey DNA was detectable significantly longer for all fragment sizes than for feces. Based on these observations, we conclude that prey DNA detected from regurgitates and whole consumers DNA extracts are comparable, whereas prey DNA detected from feces, though still sufficiently reliable for ecological studies, will not be directly comparable to the former. Therefore, regurgitates and feces constitute an interesting, non-lethal source for dietary information that could be applied to field studies in situations when invertebrate predators should not be killed.

## Introduction

DNA-based diet analysis is rapidly being employed as a widespread tool for empirically characterizing diet and trophic interactions in a broad range of vertebrates and invertebrates (Traugott *et al.* 2013; Clare 2015). DNA-based methods for diet analysis typically rely on the detection of short fragments of prey DNA, recovered from predator’s gut contents (*e.g*. Leray *et al.* 2015; Mollot *et al.* 2014) or other types of dietary samples such as feces, regurgitates, or whole consumers (*e.g*. Ibanez *et al.* 2013; Kartzinel *et al.* 2015; Thalinger *et al.* 2016; Wallinger *et al.* 2015). The success of DNA-based approaches to analyze trophic interactions is mainly due to the fact that they allow the direct and accurate identification of trophic links from minute amount of starting material, even for very small-sized organisms such as mites (Pérez-Sayas *et al.* 2015) or zooplankton (Durbin *et al.* 2012). Furthermore, the rapid growth of public sequence databases and methodological improvements in detection sensitivity and high-throughput technology, offer time- and cost-effective procedures applicable to a great variety of ecological systems and to large sample sizes (*e.g*. Valentini *et al.* 2009; Pompanon *et al.* 2012; Sint *et al.* 2011).

DNA-based diet analysis is particularly valuable for studying invertebrate generalist predators (Symondson 2012). Indeed, DNA methods offer a sensitive and flexible alternative to traditional behavioural or dissecting techniques that often fail to detect prey that does not leave hard remains in these cryptic liquid feeders (Traugott *et al.* 2013). DNA techniques are, however, also subject to bias and prey DNA detection success could be hampered by a variety of factors among which the type of dietary sample could play an important role (King *et al.* 2008; Pompanon *et al.* 2012; Traugott *et al.* 2013). In the case of arthropods, whole body extracts are usually the most convenient source of dietary DNA that avoid laborious dissections. Besides the drawbacks of a lethal approach (e.g. sacrificing rare or endangered species; affecting population dynamics, etc.), whole body extracts may pose additional challenges especially in the case of DNA metabarcoding diet analysis. As DNA metabarcoding combines general primers and high-throughput sequencing, the concomitant amplification of consumer DNA usually compromises the detection success of scarcer and degraded prey DNA (*e.g*. Shehzad *et al.* 2012; Piñol *et al.* 2014).

Waldner & Traugott (2012) demonstrated that regurgitates, a fluid mixture containing semi-digested prey remains and digestive enzymes, obtained from predatory carabid beetles provided superior prey DNA detection rates compared to whole body DNA extracts. Another prospective source of food DNA are feces, although their use as a dietary source in invertebrates is still uncommon (*e.g*. Ibanez *et al.* 2013; Redd *et al.* 2014; Sint *et al.* 2015). Usually, both regurgitates and feces seem to provide similar or better detection rates compared to whole body extracts (Durbin *et al.* 2012; Egeter *et al.* 2015; Unruh *et al.* 2016), and contain comparatively much less consumer DNA, making them putatively an ideal source for metabarcoding diet analysis. Nevertheless, to date we lack a comparative and quantitative assessment of the respective efficiency in detection success between whole bodies, regurgitates and feces as well as prospective interactions with other sources of non-dietary variation such as the target DNA fragment size.

In this study, we address this knowledge gap by comparing the prey DNA detection rates for three types of dietary samples: whole consumers including their gut content, regurgitates and feces. Using a controlled feeding experiment involving a widespread carabid predator, *Pterostichus melanarius* (Coleoptera: Carabidae) we test the following hypotheses: (i) post feeding prey DNA detection success should be similar or better in regurgitates compared to whole beetles due to lesser degradation of prey DNA in the former; (ii) prey DNA detection success should be lower in feces compared to regurgitates and whole bodies as faecal material represents the final stage of the digestion process; and (iii) prey DNA detection should decrease with increasing DNA fragment size and the time post-feeding for all types of samples.

## Material & Methods

### Sampling and maintenance of predators

*Pterostichus melanarius* individuals were collected by dry pitfall traps in two adjacent maize fields situated at the experimental site of INRA Le Rheu (Ille-et-Vilaine, France; GPS coordinates: 48.10744282N; 1.78830482W). Regular 24-hour trapping sessions were conducted in July – August 2013 until a sufficient number of individuals had been collected. All living beetles were brought to the laboratory where they were identified to species level and individually placed in plastic containers filled with loam. Beetles were stored at room temperature and continuously provided with water and food (field-collected earthworms and small pieces of apple).

### Feeding experiment

Prior to the feeding experiment, beetles were starved for 96 h in fresh individual plastic Petri dishes (5 cm diameter) containing only a droplet of water. After the starvation period, all beetles were transferred to a new Petri dish and provided with one freshly freeze-killed mealworm (*Tenebrio molitor*, Coleoptera: Tenebrionidae) cut in half. Carabids were allowed to feed for one hour in a dark climatic chamber at 20°. After feeding all beetles which had fully consumed the mealworm were transferred into fresh Petri dishes with no food. Beetles were stored at room temperature and continuously provided with water during the experiment.

For the “whole beetle” treatment, batches of 10 randomly chosen carabids were frozen in 2-mL reaction tubes by immersion in liquid nitrogen at 0, 12, 24, 36, 48, 60, 72 and 96 hours post-feeding. Immersion in liquid nitrogen was necessary as previous tests showed that living beetles do not die immediately after placement at −20°C, leading them to regurgitate into the reaction tube. After immersion, all whole beetles were stored at −20°C. Thirteen starved beetles were never allowed to feed and they were freeze-killed at 0 h to be used as negative controls. For the “regurgitate” treatment, batches of 10 randomly chosen individuals were allowed to regurgitate on a cotton wool tip according to the protocol described in Waldner & Traugott (2012) at 0, 12, 24, 36, 48, 60, 72 and 96 h post-feeding. After regurgitation, all beetles per given time-point were freeze-killed and stored at −20°C. Exactly the same procedure at each time point was applied on a control tip without touching a beetle for checking potential DNA carry-over contaminations. All samples were stored at −20°C prior to DNA extraction and PCR. For the “feces” treatment, 20 carabid beetles were placed in new Petri dishes after feeding with a droplet of clean water. Carabids were continuously checked for feces production at every 6 hours. Detected feces were immediately frozen within the Petri dish at −20°C whereupon the corresponding carabid individual was transferred into a new Petri dish. Feces production was monitored until all beetles died.

### Molecular diet analysis

Regurgitate and fecal samples were directly lysed in 200 μl TES Lysis Buffer (Macherey-Nagel, Germany) and 5 μl Proteinase K (10 mg/mL) overnight at 56°C. The whole beetles were previously ground using three 4 mm stainless steel beads (Lemoine S.A.S, Rennes, France) within a volume of 620 μl TES Lysis Buffer and 10 μl Proteinase K (10 mg/mL) per beetle. Tissues were disrupted by a 1-minute bead-beating step using a professional paint mixer (Fluid Management Inc., Wheeling, IL, USA). All samples were incubated overnight at 56°C. Respectively 2, 6, and 2 lysate blanks (i.e. no DNA material) were carried out for the whole beetles, fecal and regurgitate treatments. DNA was extracted in batches of 92 samples using the Biosprint 96 DNA Blood Kit (Qiagen, Hilden, Germany) on a Biosprint 96 extraction robotic platform (Qiagen) following the manufacturer’s instructions. DNA was finally diluted in 200 μl TE buffer (0.1 M TRIS, pH 8, 10 mM EDTA) and the extracts were stored at −28 °C. To avoid contamination, DNA extractions were done in a separate pre-PCR laboratory using a UVC-equipped laminar flow hood. To check for sample-to-sample cross-contamination, four extraction negative controls (PCR-grade RNase-free water instead of lysate) were included within each batch of 92 samples. All of these controls tested negative using the diagnostic PCR assay described below.

The DNA extracts were screened with a multiplex PCR assay targeting three overlapping COI mtDNA fragments of *T. molitor*, i.e. 128 bp, 332 bp and 612 bp. The primer mix contained 6 μM of primers Ten-mol-S210 (5’-TACCGTTATTCGTATGAGCAGTAT-3’) and Ten-mol-A212 (5’- CGCTGGGTCAAAGAAGGAT-3’) as well as 2 μM of primers Ten-mol-S232 (5’- TAATAAGAAGAATTGTAGAAAACGGG-3’) and Ten-mol-S231 (5’- TCATTTTTGGAGCGTGATCC-3’) (Oehm *et al.* 2011; Sint *et al.* 2011). Each 10 μl PCR consisted of 1.5 μl template DNA, 5.0 μl of 2x Multiplex PCR Kit reaction mix (Qiagen), 1.0 μl of primer mix, 0.5 μl of bovine serum albumin (BSA, 10 mg ml^−1^), and 2.0 μl of PCR-grade RNase-free water (Qiagen) to adjust the volume. Thermocycling was conducted in Eppendorf Mastercyclers (Eppendorf, Hamburg Germany) and cycling conditions were 15 min at 95 °C, 35 cycles of 30 sec at 94 °C, 90 sec at 63 °C, 1 min at 72 °C, and final elongation 10 min at 72 °C. To check for amplification success and DNA carry-over contamination, two positive (mealworm DNA) and two negative controls (PCR water instead of DNA) were included within each PCR, respectively.

The PCR products obtained were visualized using QIAxcel, an automated capillary electrophoresis system (Qiagen), with method AL320. The results were scored with Biocalculator Fast Analysis Software version 3.0 (Qiagen) and the threshold was set to 0.07 relative fluorescent units. Samples above this threshold and showing the expected fragment length were counted as positives. All DNA extracts that were tested negative in the first run were re-tested with general primers (Folmer *et al.* 1994) in a second PCR to check for any amplifiable DNA (all of these samples tested positive). To ensure contamination-free conditions, PCR preparation and visualization of PCR products were done in two separate laboratories (workflow: from pre- to post-PCR areas).

### Statistical analyses

A generalized linear mixed model was built to fit a logistic regression on the DNA detection data. We integrated three fixed effects into the model: two qualitative factors, the marker size (128 bp, 332 bp, 612 bp) and the sample type (regurgitates, feces or whole body DNA extracts), and one continuous variable, the time post-feeding. To compensate for non-independence in collection of feces individuals were included as a random effect. The model was fitted using the *glmm* function from the R package “*glmm*” (https://cran.r-project.org/web/packages/glmm). Models were fit using a Monte Carlo sample size of 1024 with 10,000 iterations. The distribution of each of the model parameters was approximated to a normal distribution using the maximum goodness-of-fit estimation with the “*fitdist*” function available in the R package “*fitdistrplus*” (Delignette-Muller & Dutang 2015). The variance in detectability rates explained by the model was estimated using the coefficient of determination method (Tjur 2009). Tests of the differences between mean detectability rates for each of the qualitative factors (marker length and sample type) were conducted using a Z-test. The time point for a prey detection probability of 50% (i.e. the time point at which on average half of the individuals show positive for the target prey) was determined for each dietary sample and DNA fragment size. To compensate for false discovery rate in multiple testing comparisons between fragments were based on 95% confidence limits (CI) as suggested by Greenstone et al (2013). All statistical analyses were conducted using the R software (R Core Team 2013).

## Results

Detectability of mealworm DNA in *P. melanarius* decreased with increasing post-feeding time and prey DNA fragment length for the three dietary samples (Fig. 1, small *vs* medium and small vs large fragments: *p*<0.001; medium *vs* large fragment: *p*=0.08), with post-feeding detection time intervals being longest for the shortest DNA fragment (Fig. 1 A, B, C). We also observed a significant effect of the dietary sample type, with prey DNA detection success being significantly lower in feces compared to regurgitates and whole beetles for all the three fragment sizes (Fig. 1, in all cases *p*<0.001). There was also a tendency for longer post-feeding detection periods in regurgitates compared to whole beetles (Fig. 1A, B) but differences were not significant (*p*=0.6). Our model fitted the data well for all of the three dietary samples: regurgitates (Fig. 1A), whole beetles (Fig. 1B) and feces (Fig. 1C), and explained 50% of the variance in DNA detectability. Raw data are presented in Table 1. For the small prey DNA fragment, 50% detection time was the highest for regurgitates (94 hours) but the value significantly dropped by more than half for the medium fragment (42 hours) and was significantly shortest for the largest prey DNA fragment (30.6 hours; Table 2). In feces 50% detection probabilities were the lowest for all the three DNA fragment sizes, with only 19 hours for the largest DNA fragment (612 pb) and a significantly shorter detection probability for the medium prey DNA fragment when compared to the regurgitate samples (Table 2).

**Figure 1.**
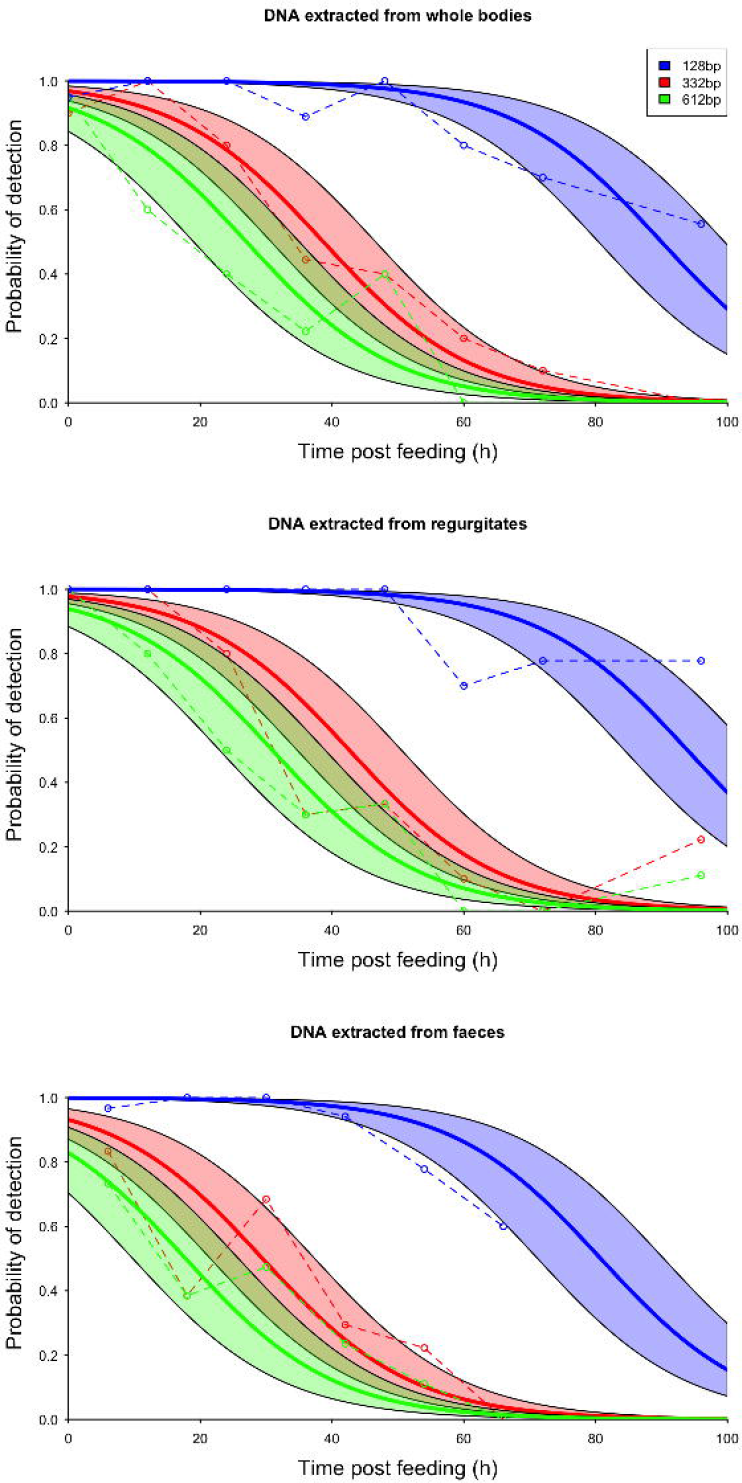
Prey DNA detection success in the predatory carabid beetle *Pterostichus melanarius* for regurgitates (A), whole bodies (B) and feces (C). Detection rates are provided for the different time points examined within each dietary sample and for the three target DNA fragment sizes. Circles and dashed lines indicate actual measures. Bold solid lines indicate the logistic regressions estimated from the glmm model and the shaded area the 95% confidence interval envelopes of the fit. The horizontal line represents the 50% prey DNA detection probability. Corresponding lower and upper 95% confidence limits are presented in Table 2.

**Figure 2.**
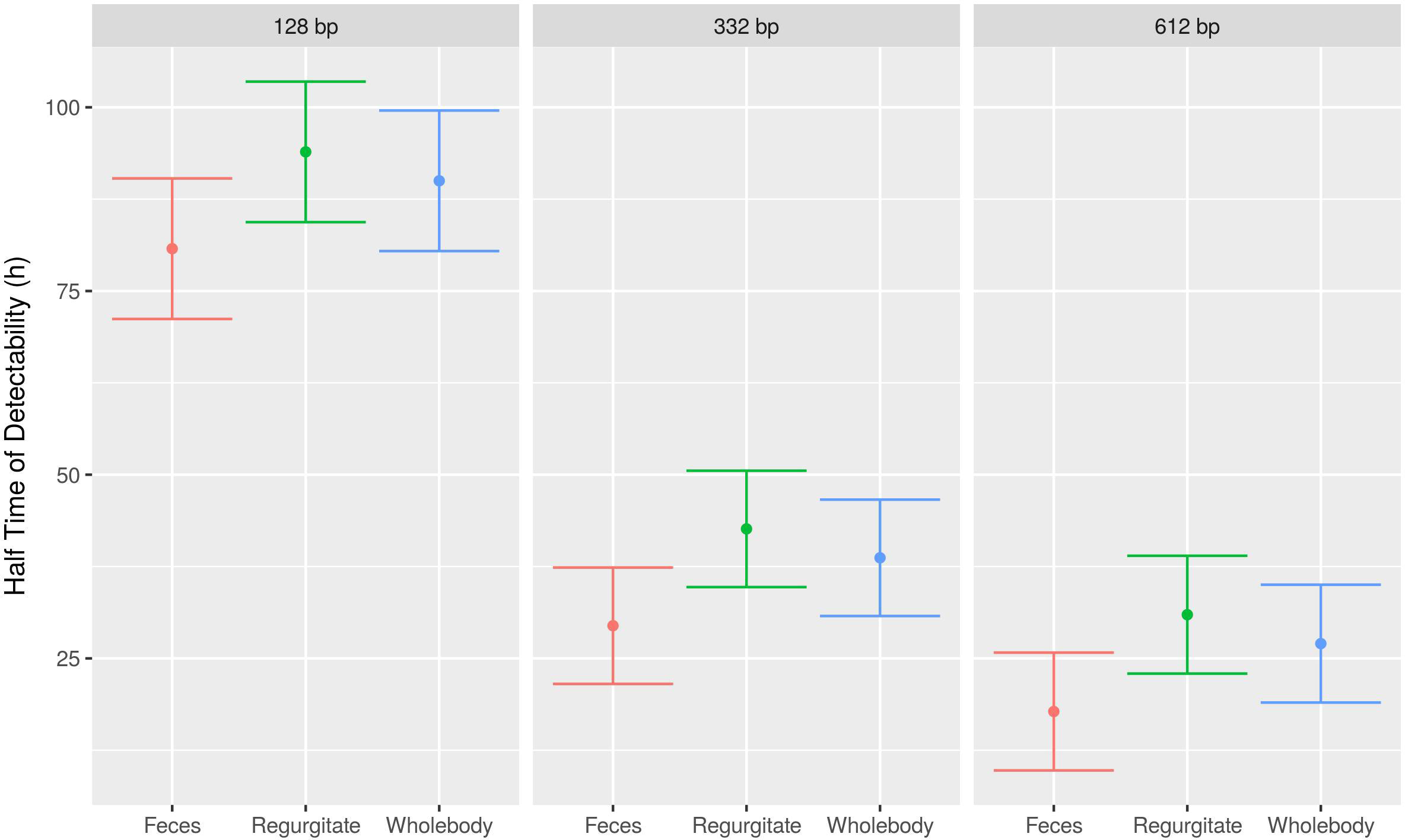
Estimated time points post-feeding for a 50% prey DNA detection probability for the different types of dietary samples and DNA fragment sizes. Provided are the 50% prey detection probabilities in hours post-feeding.

**Table 1.** Detection rates of small (128 bp), medium (332 bp) and large (612 bp) prey DNA fragments of the mealworm *Tenebrio molitor* fed to the carabid *Pterostichus melanarius* in whole beetles, regurgitates, and feces. *N* is the number of samples analyzed per digestion time.

|  |  |  | Detection rate per fragment size (%) |  |  |
| --- | --- | --- | --- | --- | --- |
| Dietary sample | Digestion time (h) | n | Small (128 bp) | Medium (332 bp) | Large (612 bp) |
| Whole bodies | 0 | 20 | 95 | 90 | 95 |
|  | 12 | 10 | 100 | 100 | 60 |
|  | 24 | 10 | 100 | 80 | 40 |
|  | 36 | 9 | 78 | 44 | 22 |
|  | 48 | 10 | 100 | 40 | 40 |
|  | 60 | 10 | 80 | 20 | 0 |
|  | 72 | 10 | 70 | 10 | 0 |
|  | 96 | 9 | 56 | 0 | 0 |
| Regurgitates | 0 | 11 | 100 | 100 | 100 |
|  | 12 | 10 | 100 | 100 | 80 |
|  | 24 | 10 | 100 | 80 | 50 |
|  | 36 | 10 | 100 | 30 | 30 |
|  | 48 | 9 | 100 | 33 | 33 |
|  | 60 | 10 | 70 | 10 | 0 |
|  | 72 | 9 | 78 | 0 | 0 |
|  | 96 | 9 | 78 | 22 | 11 |
| Feces | 3 | 8 | 100 | 87,5 | 100 |
|  | 6 | 13 | 92 | 69 | 69 |
|  | 9 | 5 | 100 | 100 | 60 |
|  | 12 | 4 | 100 | 100 | 50 |
|  | 15 | 7 | 100 | 43 | 57 |
|  | 21 | 2 | 100 | 50 | 50 |
|  | 24 | 4 | 100 | 25 | 0 |
|  | 27 | 12 | 100 | 75 | 50 |
|  | 30 | 2 | 100 | 50 | 50 |
|  | 33 | 5 | 100 | 60 | 40 |
|  | 39 | 9 | 100 | 22 | 11 |
|  | 42 | 3 | 67 | 0 | 0 |
|  | 45 | 2 | 100 | 0 | 0 |
|  | 48 | 3 | 100 | 100 | 100 |
|  | 51 | 4 | 100 | 0 | 25 |
|  | 54 | 2 | 50 | 0 | 0 |
|  | 57 | 1 | 100 | 100 | 0 |
|  | 58 | 1 | 100 | 100 | 0 |
|  | 60 | 1 | 0 | 0 | 0 |
|  | 64 | 1 | 100 | 0 | 0 |
|  | 70 | 3 | 67 | 0 | 0 |
|  | 76 | 1 | 0 | 0 | 0 |

## Discussion

How long prey DNA can be detected in a sample is determined by a range of interacting factors related to the environment, the predator-prey system and the molecular techniques used. These might affect results, but how is difficult to disentangle without conducting comprehensive experiments that explicitly account for factor multiplicity. Here, we compared multiple dietary samples from one species of invertebrate consumer, in a controlled feeding experiment, and assessed how the combined effects of the type of dietary sample and DNA fragment size will affect the prey DNA detection probability over time since feeding. Our results show that each of these factors significantly affects the rate at which the probability of detecting prey DNA decreases over time. Consistent with our hypothesis, the time during which DNA could be detected was the longest for regurgitates, for each of the three tested prey DNA fragment sizes. While this was not significantly different from DNA detected from whole beetles, prey DNA contained in feces was detectable for a significantly shorter time for all three fragment sizes.

Our results support the general assumption that regurgitates constitute a good alternative source of prey DNA (Waldner & Traugott 2012; Wallinger *et al.* 2015). Such an alternative could be particularly useful in manipulative food web experiments, where the removal and killing of the targeted predators during sampling could disturb the system under study. As 79% of predaceous land-dwelling arthropods use extra-oral digestion (Cohen 1995), this approach is potentially applicable to a large array of taxa and ecological situations. Furthermore, by containing comparatively less predator DNA, regurgitates could also be a valuable source of dietary data in DNA metabarcoding studies involving the use of general primers (Waldner & Traugott, 2012).

Nevertheless, the use of regurgitates could entail some additional limitations such as the detection of only the most recent diet items, and probably represents merely a narrow fraction of individual’s diet, especially in generalist feeders with frequent switching behavior such as carabids (Lövei & Sunderland, 1996). Thus, the choice of the most appropriate dietary sample will most likely consist in a trade-off between DNA detection rates and representativeness in terms of diet according to the focus of interest.

Fecal samples could provide a more integrated picture of individual’s diet. Our results show that overall prey DNA detection was lower compared to regurgitates and whole bodies. Note, however, that this was true only for the medium sized fragment when considering the 50% prey DNA detection probability, with significantly lower post-feeding interval found in feces compared to regurgitates. This indicates that feces overall are a good source of dietary DNA at least in *P. melanarius* beetles. Similarly, earlier study in wolf spiders showed that prey DNA was detectable in spider feces albeit in lower rates compared to whole body DNA extracts (Sint *et al.* 2015). As spiders represent an important group of generalist feeders that typically do not regurgitate, the sole non-lethal dietary sample that could be collected are feces. Also, we cannot rule out the possibility that in our case DNA prey detection success in feces was lower simply due to the constraints of the experiment. As carabids were checked for feces every 6 hours, feces deposited earlier within that timeframe could have experienced higher DNA degradation due to longer exposure to ambient temperature, thus resulting in increased variability in DNA prey detection success.

In a recent paper Unruh et al. (2016) even show that there is no difference in DNA detection between whole bodies and feces in the insect predator *Forficula auricularia*. While the authors do not discuss the possible mechanisms behind this observation, results tend to suggest that feces could be at least as good dietary source as whole body extracts for predatory insects such as *F. auricularia*. Hence, feces remain a viable non-lethal dietary source in certain situations, as detection rates are generally high.

Yet, the question of the time it takes for prey DNA to travel through the digestive tract of insects and how this varies across different taxa remains. Having a better understanding about the temporal aspects of digestion in insects in general, and particularly in carabid beetles is important. Carabids are generalist, mobile feeders with frequent switching behavior (Lövei & Sunderland, 1996) meaning that frequent diet shifts but long prey DNA retention periods may result for instance in an overestimation of consumption rates or in a mismatch between diet composition and estimations of prey availability at the place where dietary samples were collected. We also usually do not consider whether this problem could be exacerbated in herbivorous species as the digestion process of plant DNA in insects can last much longer as compared to animal DNA (Staudacher *et al.* 2011; Wallinger *et al.* 2013, 2015). For instance, results have shown that ^14^C-inulin labelled prey in carabid beetles could still be detected in feces up to five days post-feeding (Cheeseman and Gillott 1987). It will be interesting to confront these findings with observations about prey DNA transit.

Here, we also show that prey DNA detection continuously decreases over time for all the three types of dietary samples, with longer fragments (332-612 bp) decaying more rapidly compared to the shorter one (128 bp). These results meet our expectations and corroborate the general idea that digested DNA molecules break down relatively quickly and that the size of the targeted prey DNA fragment affects post-feeding prey DNA detection (Agustí *et al.* 2003; von Berg *et al.* 2008). In line with previous studies, our results support the idea that targeting short to medium size DNA fragments in DNA diet analysis is essential in order to maximize the prey detection (Deagle *et al.* 2006; Valentini *et al.* 2009). Nonetheless, if a recent feeding event is the focus, then targeting longer fragments might actually be a better strategy to ensure that only the most recent prey items are detected. Additionally, as in DNA metabarcoding diet analysis there is generally a trade-off between DNA fragment length and taxonomic resolution, targeting longer DNA fragments – within a certain range - could indeed improve the taxonomic discrimination of prey species (Pompanon *et al.* 2012). In this study, the most important observed source of variation in terms of prey DNA detection, besides time post feeding, is DNA fragment size. This could have profound implications in metabarcoding studies where the DNA fragment size usually needs to be optimized in order to meet criteria for both optimal detectability and taxonomic resolution (Taberlet *et al.* 2012). It would be interesting to simultaneously explore the decay rate of a larger array of DNA fragments of different lengths in order to assess whether a general relationship between DNA length and detectability can be drawn despite the many other sources of variability detected in previous studies. One might speculate that a consistent relationship between DNA detection success and DNA fragment size could be further used as a raw predictor of post feeding prey DNA detection intervals based solely on prey DNA fragment length.

In general, our findings show that quantitative analyses of diet based on different DNA fragment sizes or on different dietary samples are not directly comparable. Our study suggests that for estimating and comparing consumption rates for the same species between studies using different DNA fragment sizes or different dietary samples (whole beetles/regurgitates vs feces), values should be corrected after taking into account differences in detection probabilities (e.g. Greenstone *et al.* 2010). Nevertheless, prey DNA detection depends on numerous additional factors including species identity of the prey or the predator (Hosseini *et al.* 2008; Wallinger *et al.* 2013), the feeding mode (Greenstone *et al.* 2007, 2013), the time since the last meal, the number/size or the quality of prey consumed (Hoogendoorn & Heimpel 2001; Harper *et al.* 2005; Eitzinger *et al.* 2014), which we did not investigate here. The next step therefore would be the integration of multiple sources of variation in a complex multispecies, multifactorial experimental design where the different sources of variation could be quantified at once, and hierarchized (Welch *et al.* 2014).

## Acknowledgements

We thank Paul Abrams for his invaluable help with early analyses of data presented here. This study was founded by the French National Research Agency (Peerless project ANR-12-AGRO-0006). S.K. was funded by the Atlantic Canada Opportunities Agency. R.M. was supported by a grant of the regional government of Tyrol (“Land Tirol”) provided by the Mountain Agriculture Research Unit of the University of Innsbruck. We also thank the three anonymous referees for their pertinent suggestions which helped improve the present manuscript.

## Data Accessibility

All the data used in this manuscript are included in the figure and the table presented within the paper.

## Author Contributions

MT, SK and MP designed the experiment. SK realized the field work and the feeding experiment. RM carried out molecular analyses. ORR and EC realized data analyses. SK wrote the manuscript with input from all the authors.

## Conflicts of Interest

The authors have declared that no competing interests exist.

